# Metabolic rewiring of sugar alcohols drives lichen symbiosis in Trebouxiophyceae microalgae

**DOI:** 10.64898/2026.11.13.766367

**Authors:** Marta Pérez-Rodrigo, Daniel Calvo, Pedro Carrasco, Francisco Marco

**Author notes:** Corresponding author: Marta Pérez-Rodrigo. Department of Biochemistry and Molecular Biology, Faculty of Biology, University of Valencia - BIOTECMED. Address: C/ Dr. Moliner, 50, 46100 Burjassot (Valencia, Spain).

## Abstract

- Lichen symbioses are among the most successful mutualisms in extreme environments, yet the biochemical traits that enable algal partners to thrive within them remain poorly defined. We used untargeted GC-MS to uncover metabolic rules underlying symbiotic compatibility. Comparing 21 Trebouxiophyceae species, including free-living microalgae and lichen photobionts, we identified 260 metabolites, concluding that sugar alcohol metabolism emerged as the principal axis of differentiation, with ribitol-rich taxa consistently dominating lichen partnerships. High ribitol producers shared a coordinated metabolic program coupling elevated malate with compatible solutes (glycerol, inositol, proline) and phospholipid intermediates, supporting osmoprotection, redox balance, and membrane remodeling.
- They released ribitol into the culture medium together with a distinctive suite of metabolites (including maltose, allulose, and tyramine) not previously reported in lichen algae, suggesting new candidates for symbiotic signaling.
- A comparative analysis of *Ramalina farinacea* and its photobiont, *Trebouxia jamesii*, revealed a metabolic shift in symbiosis towards sustained arabitol, ribitol, and sucrose accumulation, and away from glucose accumulation, indicating an adaptive carbon-allocation strategy.
- These findings identify ribitol-centered metabolic rewiring as a pivotal trait for lichenization, provide mechanistic insight into stress resilience and partner specificity, and establish metabolomic profiling as a predictive framework for chemotaxonomy and testing algal-fungal compatibility in symbiotic systems.

## Introduction

Microalgae are widely recognized for their ability to produce a wide array of bioactive compounds with significant biotechnological potential. These metabolites have valuable applications across various fields, including medicine, cosmetics, and food science, due to their antioxidant, antibacterial, and anticancer properties (Levasseur *et al*., 2020; Ibrahim *et al*., 2023). Beyond their practical uses, microalgae play a vital ecological role as primary producers, contributing to oxygen generation and nutrient cycling in both aquatic and aero-terrestrial ecosystems (Metting, 1996; Leliaert *et al*., 2012). Their adaptability to diverse environments also makes them key contributors to ecosystem resilience and stability, influencing factors like soil health and water quality (Nava & Leoni, 2021; Joseph & Ray, 2024).

This study centers on microalgae of the class Trebouxiophyceae, a diverse group of unicellular green algae within the phylum Chlorophyta. Among its members, the genus *Chlorella* is especially well-known and widely used in research and industry due to its easy cultivation (Krienitz *et al*., 2015). Trebouxiophyceae species occur both free-living and as symbionts, most notably in lichens (Veselá *et al*., 2024). This class is of particular interest because it comprises the largest group of green algal photobionts, accounting for most of the approximately 150 Chlorophyta species currently known to engage in lichen symbioses (Sanders & Masumoto, 2021). Photobionts are photosynthetic microorganisms, green algae (phycobionts) or cyanobacteria (cyanobionts), that form symbiotic associations with a heterotrophic fungal partner (mycobiont) in lichens, often together with additional microorganisms (Grimm *et al*., 2021).

Microalgae capable of forming symbiotic relationships are also found in free-living states in nature (Tschermak-Woess, 1978; Bubrick *et al*., 1984; Spribille *et al*., 2022), which indicates that they have the metabolic capability to thrive in both lifestyles. However, the specific traits that make these algae suitable for lichenization, as well as the metabolic mechanisms underlying their compatibility with their specific lichen partners, remain poorly understood.

In addition, lichens inhabit harsh, highly variable, and restrictive habitats. They thrive in ecological niches where most vascular plants cannot survive, such as arid deserts, polar regions, seashores, high-altitude mountains, and other extreme environments. Their wide distribution across diverse biomes highlights their ecological importance and adaptability, emphasizing the need to study the biochemical processes that shape their stress tolerance (Lorenz *et al*., 2023; Canali *et al*., 2025).

Since the initial studies on the biochemical components of lichen symbiosis in the 1960s, it has been shown that one of the main roles of the photobiont in the lichen is to provide the fungal partner with photosynthesis-derived products, mainly sugars and sugar alcohols, which are released extracellularly and support the mutualistic relationship and resilience of the symbiosis (Drew & Smith, 1967; Richardson *et al*., 1967). Research on the compounds produced by these microalgae has traditionally focused on these molecules, linking them to their role in symbiosis and/or abiotic stress tolerance (Kranner *et al*., 2022), while the comprehensive metabolomics of these microalgae remains unexplored.

Within the family Trebouxiaceae, microalgae from the order Trebouxiales participate in a highly diverse number of lichen symbioses and represent the main photobiont species (Palmqvist *et al*., 2008), while they can also occur as free-living organisms (Veselá *et al*., 2024). Early studies showed that species of *Trebouxia, Asterochloris*, and *Myrmecia* isolated from different lichens release ribitol (Richardson *et al*., 1968; Komiya & Shibata, 1971; Hill & Smith, 1972; Lines *et al*., 1989). More recent analyses have further confirmed ribitol production by *Trebouxia* (Kosugi *et al*., 2013; Sadowsky *et al*., 2016). However, there are no reports on compounds produced in other Trebouxiales genera within this clade, such as *Symbiochloris* or *Vulcanochloris*.

Another order within Trebouxiophyceae is Chlorellales, which belongs to the family Chlorellaceae. A notable representative from this clade is *Chlorella vulgaris*, which is not known to form lichen symbioses, but it does engage in symbiotic relationships with protists and cnidarians. Metabolomic analyses of *C. vulgaris* reveal a distinctive metabolic profile, producing compounds such as sugars, fatty acids, and notably low levels of the sugar alcohol myo-inositol. The trace detection of sugar alcohols suggests that disaccharides predominantly shape the sugar profile of this microalga (Gustavs et al., 2011; Zhang et al., 2014; Qu et al., 2021; Hegazi et al., 2024). Other Chlorellales, such as *Coenochloris* (family Oocystaceae), remain largely unexplored at the metabolic level. These microalgae are typically found as free-living organisms in biofilms (Komar *et al*., 2023).

Another order within this group is Prasiolales, which includes the family Stichococcaceae. Within this clade, studies on the genus *Diplosphaera*, specifically on the species *Diplosphaera chodatii*, have shown that it releases high quantities of sorbitol, together with glucose and sucrose (Wang *et al*., 2014; Medwed *et al*., 2021; Nazem-Bokaee *et al*., 2025). *Diplosphaera* occurs both as a free-living organism (Medwed *et al*., 2021) and as a photobiont in lichen symbioses (Fontaine *et al*., 2013; Doering *et al*., 2020; Chiva *et al*., 2023). It has been observed that, in some cases, these algal cells are smaller than those in the main thallus and are not lichenized (Sanders & Masumoto, 2021).

*Coccomyxa* green algae belong to the family Coccomyxaceae, within the class Trebouxiophyceae. Some studies report how *Coccomyxa* species produce variable quantities of ribitol (Hill & Smith, 1972; Gustavs et al., 2011), whereas other species like *C. melkonianii* or *Coccomyxa* sp. IITRSTKM4 are not described to produce detectable levels of this compound (Fais *et al*., 2021; Tripathi *et al*., 2021). The habitats of the *Coccomyxa* species are very diverse, with some species found as free-living organisms in terrestrial and aquatic environments, and others in lichen symbiosis or symbiosis with other organisms. Interestingly, the cell walls of *Coccomyxa* photobionts are typically not penetrated by the mycobiont in lichen symbiosis (Sanders & Masumoto, 2021).

Other genera, such as the *Watanabea* from the order Watanabeales (family Watanabeaceae), remain largely unexplored in terms of metabolite production. Within this genus, species like *W. acidophila* are usually found as free-living organisms, but not exclusively. This species, together with *W. lichenicola,* discovered in lichen *Buellia zoharyi* (Chiva *et al*., 2021) has an associated (not primary) phycobiont role in lichen symbiosis.

Beyond the genera noted above, other studies targeting the detection of sugar alcohols report how several green microalgae taxa produce tetritols, pentitols, and, more rarely, hexitols, including algae described as lichen photobionts and others described as free-living aeroterrestrial green algae. The main reported compounds are erythritol, glycerol, ribitol, arabitol, sorbitol, and volemitol, in genera like *Apatococcus, Chloroidium, Prasiola, Rosenvingiella, Stichococcus*, *Trentepohlia,* and others (Lewis & Smith, 1967; Feige & Kremer, 1980; Jacob et al., 1991; Honegger et al., 1993; Gustavs et al., 2010, 2011). However, cyanobacterial photobionts primarily export glucose (Drew & Smith, 1967; Spribille *et al*., 2022).

This study aims to expand current knowledge of the compounds produced and exported by Trebouxiophyceae using modern, untargeted metabolomic techniques. It broadens the focus beyond sugars and sugar alcohols to include intermediate metabolites involved in central metabolic pathways, as well as key compounds such as amino acids, fatty acids, sterols, and secondary metabolites.

Analyses were conducted on algae both in isolated cultures and in their natural role as phycobionts within lichens under real environmental conditions (without separating the partners). The lichen *Ramalina farinacea* was chosen as the model system due to its ecological importance and well-documented symbiotic relationship (Molins *et al*., 2021).

The main objective of this work was to characterize the intracellular and excreted metabolomes of representative Trebouxiophyceae microalgae and the lichen *Ramalina farinacea*, to relate metabolite profiles to phylogenetic lineage and ecological roles (including phycobiont status and extremotolerance), identify candidate signaling molecules for mycobiont-phycobiont recognition, and determine which metabolic differences underlie the ability of certain algae to form lichen symbioses.

## Materials and methods

### Biological materials

Green microalgae used in this study were obtained from the Symbiotic Algal collection of the University of Valencia (ASUV, 2025). The twenty-one Trebouxiophyceae species selected belong to the genus *Asterochloris*, *Chlorella*, *Coccomyxa, Coenochloris, Diplosphaera, Myrmecia, Trebouxia* (clades A, C, and I), *Vulcanochloris,* and *Watanabea*. Information about strain identity is reported in Table 1. Lichen thalli samples of *Ramalina farinacea* were collected from El Toro, Castellón, Spain.

**Table 1.** Microalgal species analyzed for metabolite production using Gas Chromatography-Mass Spectrometry (GC-MS).

| **Microalgae species** | **Species authorities** | **ASUV Strain ID** |
|---|---|---|
| *Asterochloris erici* | (Ahmadjian) Skaloud & Peksa, 2010 | ASUV 53 |
| *Asterochloris glomerata* | (Warén) Skaloud & Peksa, 2010 | ASUV 56 |
| *Chlorella vulgaris* | Beyerinck [Beijerinck], 1890 | ASUV 134 |
| *Coccomyxa arvernensis* | Jaag, 1933 | ASUV 7 |
| *Coenochloris* sp. |  | ASUV 148 |
| *Diplosphaera chodatii* | Bialosuknia, 1909 | ASUV 155 |
| *Myrmecia israelensis* | (S. Chantanachat & H. Bold) T. Friedl, 1995 | ASUV 109 |
| *Symbiochloris symbiontica* | (Tschermak-Woess) Skaloud, Friedl, A.Beck & Del Grande, 2016 | ASUV 157 |
| *Trebouxia asymmetrica* (Clade A) | Friedl & Gärtner, 1988 | ASUV 128 |
| *Trebouxia corticola* (Clade C) | (P. A. Archibald) Gärtner, 1985 | ASUV 80 |
| *Trebouxia cretacea* (Clade A) | Voytsekhovich & Beck, 2015 | ASUV 133 |
| *Trebouxia decolorans* (Clade A) | Ahmadjian, 1960 | ASUV 41 |
| *Trebouxia impressa* (Clade I) | Ahmadjian, 1960 | ASUV 52 |
| *Trebouxia jamesii* (Clade A) | (Hildreth & Ahmadjian) Gärtner, 1985 | ASUV 129 |
| *Trebouxia lynniae* (Clade A) | Barreno, 2022 | ASUV 44 |
| *Trebouxia vagua* (Clade A) | Voytsekhovich & Beck, 2015 | ASUV 146 |
| *Trebouxia* sp. *arnoldoi* (Clade A) |  | ASUV 137 |
| *Trebouxia* sp. OTU A25 (Clade A) |  | ASUV 138 |
| *Vulcanochloris guanchorum* | Vancurová, Peksa, Nemcová & Skaloud, 2015 | ASUV 139 |
| *Watanabea acidophila* | Lukesová, Darienko & Pröschold, 2019 | ASUV 141 |
| *Watanabea lichenicola* | Chiva, Dumitru, Bordenave & Barreno, 2021 | ASUV 142 |

### Culture conditions and sampling

Microalgae axenic cultures were propagated in solid 3N-BBM-GC (3×N Bold’s Basal Medium (Bischoff & Bold, 1963) supplemented with glucose (20 g l^−1^) and casein hydrolysate (10 g l^−1^). Cultures were maintained in a controlled environment incubator at 20 ± 1 °C and illuminated at 25 μmol photons m^−2^ s^−1^ PAR with a 12:12 h light-dark photoperiod. The 21-day-old precultures were suspended in liquid 3N-BBM-GC medium, homogenized using a vortex, and filtered through a 100 μm pore diameter filter. A volume of 50 μl of the evenly suspended microalgae (5×10^5^ cells approx.) was inoculated on sterile cellulose-acetate filter paper discs (13 mm diam., 0.44 μm pore size) placed on solid 3N-BBM-GC. Samples were collected after 21 days of growth.

Additionally, selected microalgae were cultured in liquid 3N-BBM-GC to get liquid growth media samples. The 21-day-old precultures of *A. erici*, *T. jamesii*, *W. lichenicola,* and *C. vulgaris* were inoculated in 25 ml of 3N-BBM-GC liquid medium at an initial concentration of 1×10^6^ cells/ml in 100 ml flasks in an orbital shaker at 120 rpm with the same temperature and light conditions as plate culture. Culture media samples were collected in the late exponential phase of growth.

All the collected samples were immediately frozen in liquid nitrogen and stored at -80 °C. Before metabolite extraction, the samples were lyophilized overnight.

Lichen thalli samples of *Ramalina farinacea* were collected from *Pinus sylvestris* bark in spring and lyophilized overnight. The molecular analysis determined their primary phycobiont as *T. jamesii* (Moya *et al*., 2024).

### Phylogenetic analysis

The identity of the samples was verified by sequencing the nuclear ribosomal ITS fragment (nrITS) with subsamples of the precultures. Total genomic DNA was extracted with the DNeasy Plant Mini kit (Qiagen) following the manufacturer’s instructions (QIAGEN, http://www.qiagen.com). Algal nrITS DNA was amplified using the primers nr-SSU-1780 and ITS4 and fungal nrITS DNA using the primers ITS1F and ITS4a according to the protocol described by (Molins *et al*., 2018) PCR products were purified with the NZYGelpure kit (Nzytech, https://www.nzytech.com), and the resulting DNA samples were sequenced by Sanger and BLASTn searched against the database.

A multiple sequence alignment (MSA) for the algal nrITS was performed using MAFFT v7.526 (Katoh *et al*., 2009) and the phylogenetic tree was constructed using the Neighbor-Joining method, computing evolutionary distances based on the Jukes-Cantor model. The phylogenetic visualization was performed with iTOL v6.9 (Ciccarelli *et al*., 2006).

### Metabolome determination by GC-MS

First, 15 mg dry weight (DW) of algal biomass per species was weighed on a precision balance. The algal material was finely ground using approx. 300 mg of glass beads (diam. 500-750 μm) with a vortex with a Vortex-Genie 2 vortex attachment at maximum power for 15 minutes.

The liquid-phase extraction of metabolites and the chemical derivatization were carried out according to the protocol reported by (Lisec *et al*., 2006) with major modifications. A volume of 1.4 ml of HPLC grade methanol (pre-cooled to -20 °C) was added to the grounded sample along with 350 μl of antipyrine stock solution (10 mM in methanol) as internal standard (IS). The mixture was incubated at 70 °C, shaking at 950 rpm for 10 min, then centrifuged for 10 min at 11000 g, and 1.2 ml of the supernatant were transferred to a 1.5 ml tube with PCR clean purity grade. Aliquots (400 µL each) of each extract were dried in a vacuum concentrator for subsequent chemical derivatization.

Derivatization consisted of methoxyamination and subsequent silylation. The derivatization process was also carried out on an empty tube as a blank control. For methoxyamination, 40 µl of methoxyamine hydrochloride (20 mg/ml in pyridine) were added. This reagent was prepared just before the performance of each experiment. The samples were incubated at 650 rpm agitation for 2 h at 37 °C. For silylation, 70 μl of the N-Methyl-N-(trimethylsilyl) trifluoroacetamide (MSTFA) reagent were added to each sample, and 5 µl of C7 - C30 saturated alkanes to the first and last measured samples of each batch to calibrate retention indexes. Samples were incubated at 650 rpm for 30 min at 37 °C. Derivatized aliquots were transferred to glass insert vials for GC-MS analysis.

Gas Chromatography coupled to Mass Spectrometry (GC-MS) analysis was performed with an Agilent 5977A mass spectrometer with a quadrupole analyzer for capillary columns (split/splitless, pulsed split, and pulsed splitless). Helium was used as the carrier gas, with a constant 1 ml/min flow. The injection mode was splitless: 79.6 ml/min for 1 min. An apolar HP-5MS UI 30 m - 0.25 mm - 0.25 μm capillary column (19091S-433UI, Agilent) was used. The following temperature program was applied: the initial temperature was 60 °C for 5 min, followed by a ramp of 10 °C/min to 250 °C for 5 min and a second ramp of 20 °C/min to 300 °C for 10 min. The transfer line temperature was kept at 280 °C. The temperature of the MS source was 230 °C, and that of the quadrupole was 150 °C. The sample injection volume was 2 µl. Mass spectra were acquired at 70 eV by electron impact with a scanning range of 30 to 650 amu. The solvent delay was 8 minutes.

The system has integrated peak deconvolution and spectral matching for target identification and quantification in a complex array. System data acquisition software was MassHunter GC/MS Acquisition, Agilent Technologies. Inc. The selective mass detector operated in full scan or Scan mode, which consists of scanning between two masses to obtain total information on the content of the sample to be analyzed. The areas of the peaks corresponding to each of the compounds were integrated using the MassHunter Qualitative Software. The metabolites were identified according to the coincidence of their mass spectra against the commercial library of NIST (National Institute of Standards and Technology, Gaithersburg, MD, USA) (Ausloos *et al*., 1999). Subsequently, mass spectral matches of metabolites of interest were manually analyzed for each against the Golm Metabolome Database (GMD) mass spectral database (Kopka, 2006). Abundance data were normalized to the IS and the sample dry weight. Pure standards were used to confirm the identity and quantification of some of the compounds of interest.

### Statistical analysis of metabolite data

Metabolite data were processed and analyzed using R software v. 4.5. (R Core Team, 2025). Principal component analysis (PCA) was conducted using the factoextra package (v.1.0.7) (Package "factoextra", Kassambara & Mundt, 2025) to identify overall patterns in the datasets and visualize clustering among microalgae species based on metabolite profiles. Differences in metabolite levels among individual species were assessed through one-way ANOVA, complemented by post-hoc comparisons implemented with the multcompView package (v. 0.1-10) (Package “multcompView” Visualizations of Paired Comparisons, 2025). To evaluate differences between defined groups of species, linear mixed models (LMM) were applied using the lme4 (v. 1.1-37) and lmerTest (v. 3.1-3) packages (Kuznetsova *et al*., 2017; *Package ‘lme4’*, 2025), which allowed the inclusion of random effects to account for species-related variability. Intersections and shared metabolite composition among species were visualized using UpSet and Venn diagrams generated with the Intervene web server (Khan & Mathelier, 2017) and the Venny web tool (v. 2.1) (Venny; Oliveros, 2007), respectively. Additionally, heatmaps and similarity matrices were constructed using the Morpheus web server (Morpheus, https://software.broadinstitute.org/morpheus), with the most represented differential metabolites determined by ANOVA. Finally, metabolic pathway mapping was performed using annotated pathways derived from the KEGG pathway database (Kanehisa *et al*., 2025), which provided a functional context for the identified metabolites.

## Results

### Metabolome diversity among microalgae

Metabolite profiles of the selected microalgae (Table 1) were obtained via GC-MS analysis of methanol extracts from cultures grown on solid media for 21 days. A total of 260 metabolites were identified (Supporting Information Table S1). The highest metabolite diversity was observed in *Watanabea acidophila*, with 85 distinct compounds, whereas *Symbiochloris symbiontica* presented the lowest diversity, with only 40 metabolites identified. Four metabolites — glucose, maltose, ribitol, and sucrose — were consistently detected across all analyzed species (Figure 1a; Supporting Information Table S2). In terms of unique metabolites, *Asterochloris erici* displayed the highest number, with 12 exclusive compounds, followed closely by *S. symbiontica* and *Trebouxia* sp. *arnoldoi*, each with 11 unique metabolites.

**Figure 1.**
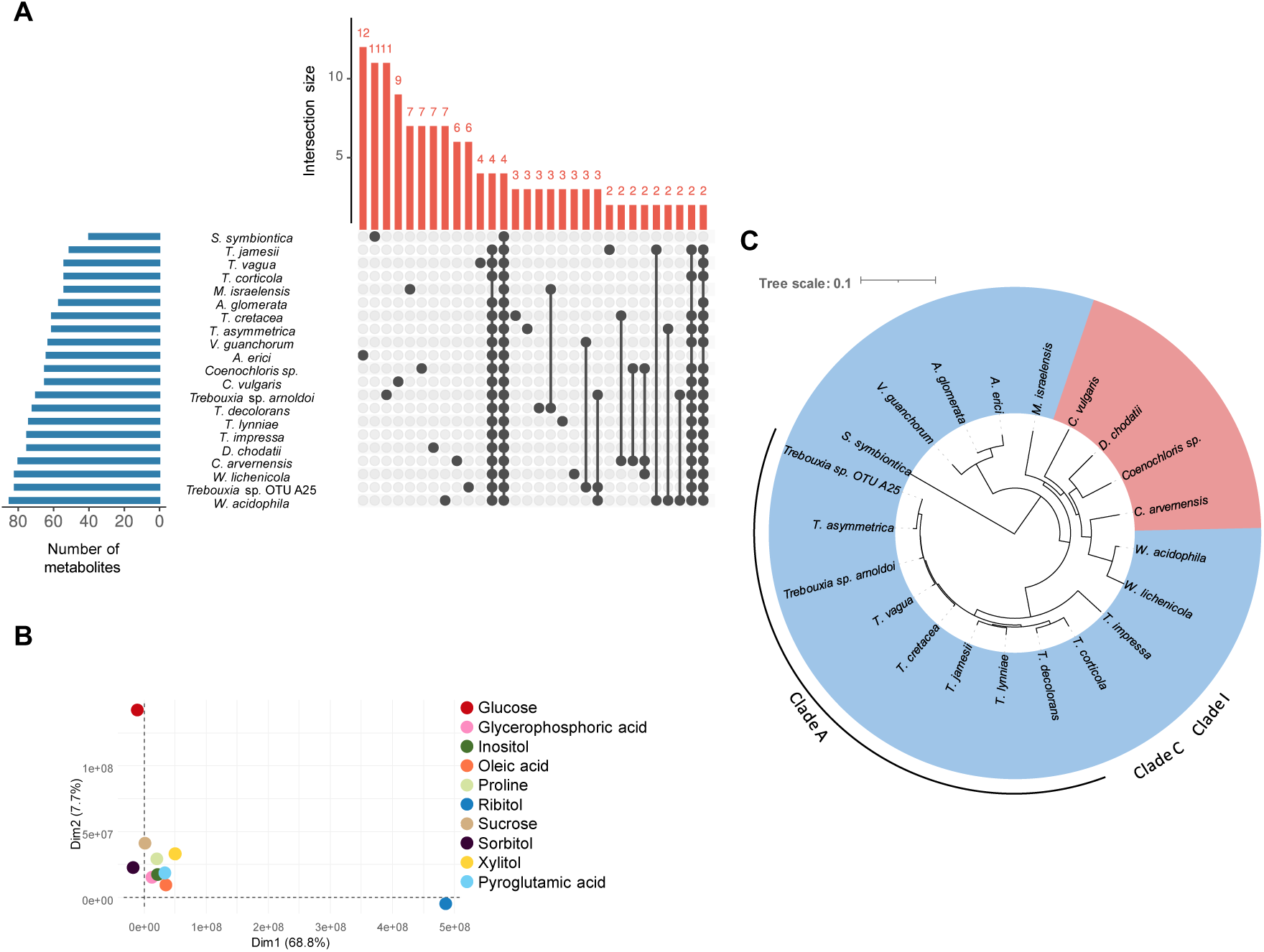
Comparative analysis of metabolite profiles across 21 species of green microalgae. a) UpSet diagram showing the distribution and overlap of identified metabolites among the analyzed species. The upper bar plot represents the number of unique and shared compounds represented with connected dots. Bars on the left indicate the total number of metabolites detected in each species. b) Principal Component Analysis (PCA) representing the 10 metabolites with higher significant variations between microalgae. The two principal dimensions are determined by ribitol and glucose. c) Phylogenetic tree of the 21 studied microalgae species based on ITS multiple sequence alignment and Neighbor-Joining phylogeny. Microalgae with high levels of ribitol are marked in blue and with low levels in red.

Each species exhibited a distinct metabolite profile (Supporting Information Figs. S1 and S2; Supporting Information Table S3). PCA comparison of the metabolome profiles revealed that several compounds contributed significantly to interspecies variations (Figure 1b), with ribitol and glucose being the primary drivers of the two principal components, followed by xylitol, sucrose, and oleic acid. Specifically, ribitol was the predominant compound in 17 of the microalgal species, whereas sucrose dominated in *C. arvernensis* and *D. chodatii*, and glucose was most abundant in *C. vulgaris* and *Coenochloris* sp. (Supporting Information Fig. S2). Interestingly, the predominance of ribitol overlaps with the phylogenetic tree distribution of the microalgae, except for *W. acidophila* and *W. lichenicola* (Figure 1c). A similar pattern was observed in the chemotaxonomic distribution of the species, based on the complete metabolomic profiles (Supporting Information Fig. S3). This observation was further supported by the PCA of the complete metabolomes, where microalgae species with high and low ribitol production are separated along the first principal component, which accounted for 68,8% of the total variation (Supporting Information Fig. S1).

Ribitol, glucose, and sucrose were among the most frequently occurring compounds within the set of 48 more represented metabolites that showed significant variations among microalgae. This group also included 18 additional sugars and sugar alcohols. Other metabolite categories represented in this set were twelve amino acids, eight organic acids, two fatty acids, and a monoacylglycerol derivative, two sterols, α-tocopherol, and phosphoric acid (Figure 2).

**Figure 2.**
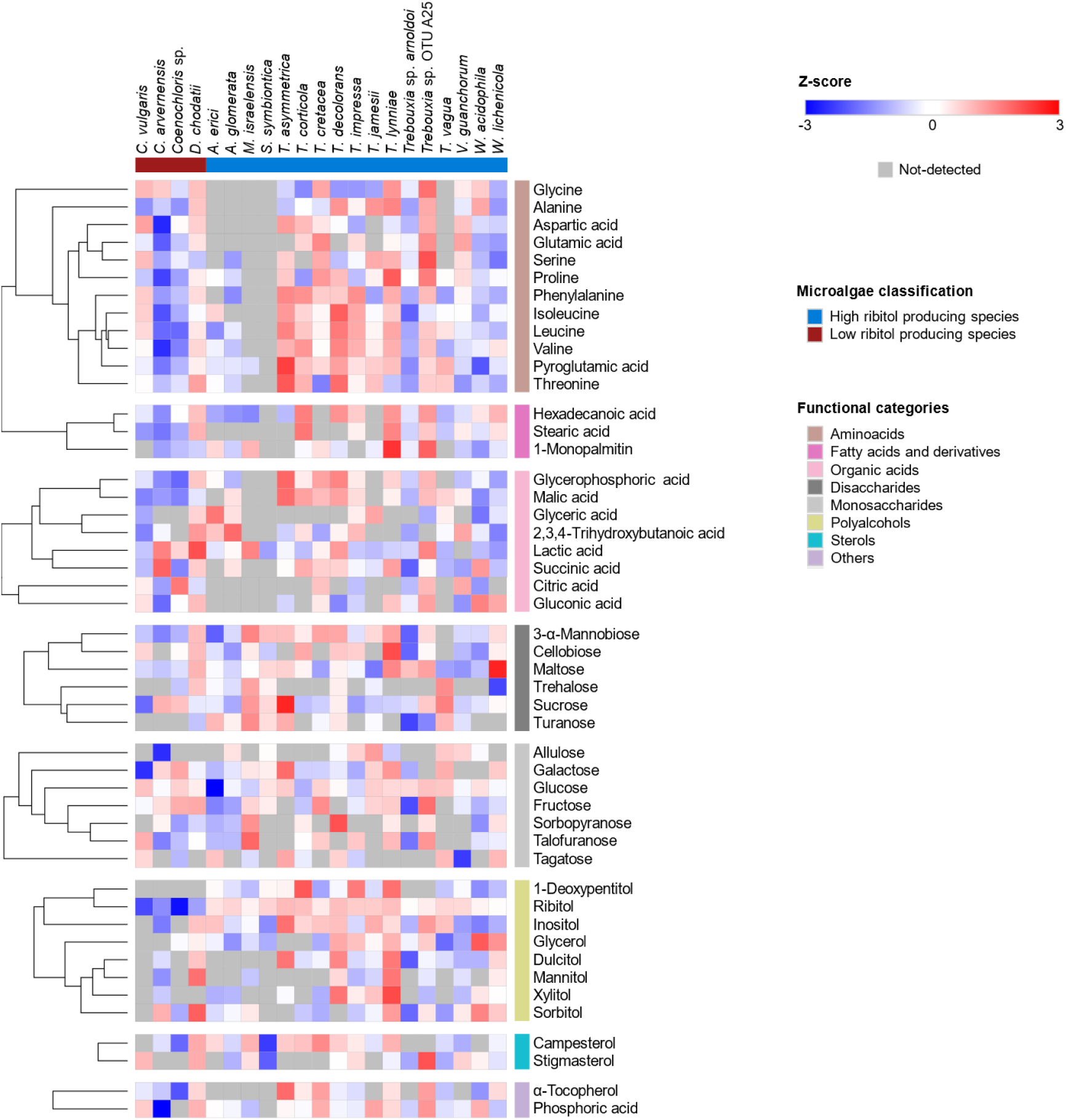
Hierarchical clustering heatmap of the 48 most represented significant metabolites across species, grouped by functional categories. Clustering was based on the metric "One minus Pearson correlation" with the average linkage method. A one-way ANOVA was performed for each metabolite to determine statistical significance (p-value < 0.05). Data were log_2_-transformed, and z-scores were calculated for each metabolite.

The pentitol ribitol was the only sugar alcohol detected across all analyzed species (Figure 2; Supporting Information Table S4). As previously noted, ribitol was the most abundant compound in *Asterochloris*, *Myrmecia*, *Symbiochloris*, *Trebouxia*, *Vulcanochloris*, and *Watanabea* species. In contrast, *Chlorella*, *Coenochloris*, *Diplosphaera*, and *Coccomyxa* exhibited only low levels of this polyol. Notably, *Trebouxia asymmetrica* showed the highest content in ribitol. Another widely distributed sugar alcohol was the cyclic hexitol inositol, which was the second most abundant polyalcohol in all *Trebouxia* species except for *T. lynniae*. *T. asymmetrica* also showed the highest inositol content, followed by *Trebouxia* sp. OTU A25, *T. jamesii*, *T. decolorans*, *T. vagua, Diplosphaera chodatii*, *Asterochloris erici*, and *T. cretacea.* Among all species, *T. lynniae* showed the highest total sugar alcohol content, whereas *C. vulgaris* showed both the lowest levels and the least diversity of these compounds. By contrast, in *D. chodatii*, and *C. arvernensis* the most abundant polyol was the hexitol sorbitol, with *D. chodatii* producing the highest levels of this compound among all the studied species. In *Coenochloris* sp., the most abundant polyol was the triol glycerol, although it did not dominate in the overall metabolome. Additionally, each species exhibited a unique sugar alcohol profile: *Watanabea* species had the highest glycerol content, *Symbiochloris symbiontica* was richest in threitol, *T. asymmetrica* in dulcitol, *T. lynniae* in xylitol, *T. corticola* in 1-deoxypentitol, and *D. chodatii* in mannitol.

A wide range of non-alcoholic sugars was also identified across the analyzed species (Figure 2; Supporting Information Table S5). Among these, *T. asymmetrica* exhibited the highest total monosaccharide abundance. Glucose was consistently the most abundant monosaccharide accumulated across all species except for *A. erici*. Notably, *Chlorella vulgaris* had the highest glucose content, representing the most abundant compound detected in its metabolome, as previously highlighted. Monosaccharides such as fructose displayed significant variability among species, reaching their highest abundance in *Trebouxia* sp. OTU A25. In contrast, galactose was predominant in *T. asymmetrica*, whereas ribose was particularly abundant in *M. israelensis*, and allulose in *T. jamesii*. Sucrose was highly abundant across the analyzed microalgae, emerging as the predominant disaccharide in all species. Indeed, it represented the major metabolite detected in the metabolomes of *D. chodatii* and *C. arvernensis*, with the highest overall content observed in *T. asymmetrica*. Another disaccharide, maltose, was also consistently present in all species and was notably enriched in *W. lichenicola*. The disaccharide 3-α-mannobiose was detected in most species and was particularly abundant in *M. israelensis*. Moreover, *T. lynniae* was distinctive for exhibiting the highest levels of cellobiose.

Differential levels of free amino acids were detected across the analyzed species (Figure 2; Supporting Information Table S6), although most amino acids remained below the detection limit in *S. symbiontica* and *M. israelensis*. *T. asymmetrica* exhibited the highest total abundance of free amino acids, followed closely by *T. lynniae.* The overall amino acid profiles showed strong phylogenetic differentiation. All species except for *S. symbiontica* accumulated pyroglutamic acid (5-oxoproline), a non-proteinogenic amino acid and intermediate of the glutathione cycle. This compound was the most abundant free amino acid in *T. vagua*, *T. decolorans*, *T. asymmetrica*, *T. corticola*, *T. impressa*, *W. lichenicola*, and *A. glomerata*. In contrast, proline was preferentially accumulated in *A. erici*, *Trebouxia* sp. OTU A25, *T. cretacea*, and *T. lynniae*. Some exceptions were observed among the remaining high ribitol producing species, *T. jamesii* and *W. acidophila* exhibited higher levels of alanine, while *Trebouxia sp. arnoldoi* and *V. guanchorum* accumulated more glutamic acid. Meanwhile, the non-proteogenic pipecolic acid emerged as the dominant amino acid only in *M. israelensis*. Interestingly, in the species with low ribitol production *Coenochloris* sp., *C. arvernensis*, and *D. chodatii*, the most abundant amino acid was 4-aminobutanoic acid (GABA), a non-proteinogenic gamma-amino acid (Figure 3). Notably, *C. vulgaris* diverged from this trend, showing a distinct profile dominated by aspartic acid.

**Figure 3.**
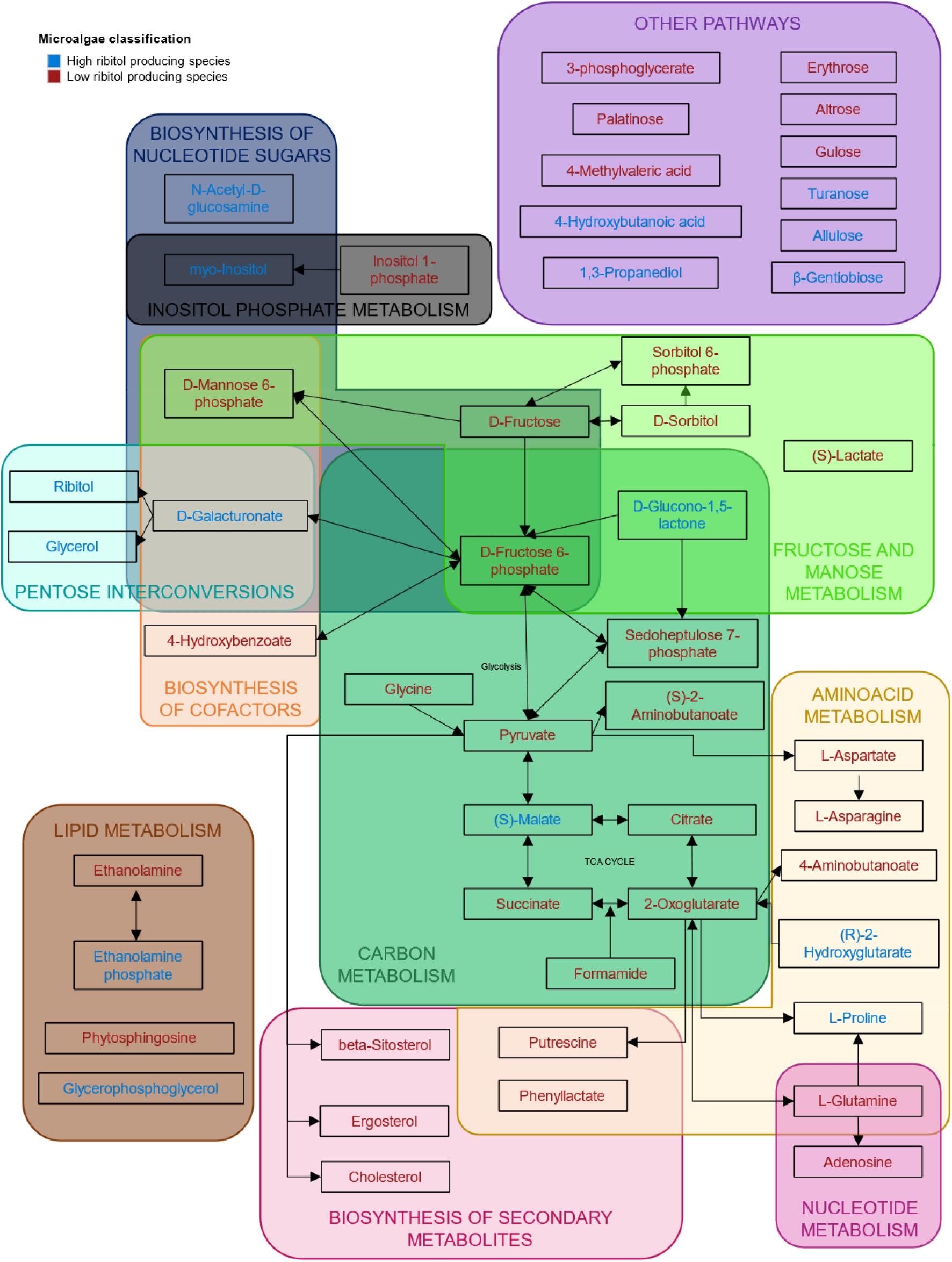
Metabolic map of differentially expressed metabolites between high-ribitol (blue) and low-ribitol (red) producing species. The pathways supported by KEGG (KEGG pathway) are included, compounds not established in KEGG pathways are included in others. A one-way ANOVA was performed to determine statistical significance (p-value < 0.05).

The content of fatty acids and their derivatives also varied considerably among the analyzed algae (Figure 2; Supporting Information Table S7). *Trebouxia lynniae* exhibited the highest total abundance of these compounds, closely followed by *T. corticola*, while *T. asymmetrica* showed the lowest levels. Oleic acid emerged as the most abundant fatty acid in eight of the analyzed species, including *Trebouxia* sp. OTU A25, *T. lynniae*, *T. corticola*, *T. impressa*, *Coenochloris* sp., *Diplosphaera chodatii*, *M. israelensis* and *W. lichenicola*. Despite its widespread presence, the levels of this monounsaturated fatty acid did not differ significantly between species. A similar trend was observed for its polyunsaturated derivative, linoleic acid, which was the predominant fatty acid in *Trebouxia sp. arnoldoi*, *C. vulgaris* and *Symbiochloris symbiontica*. In contrast, hexadecanoic acid (palmitic acid), a saturated fatty acid, was the most abundant in *T. decolorans, W. acidophila* and *C. arvernensis*, while stearic acid stood out as the main fatty acid in *T. vagua* and *V. guanchorum*. Both hexadecanoic and stearic acids were produced in higher quantities by *T. corticola.* Notably, the uncommon and rarely reported 5,8,11-eicosatriynoic acid was the most abundant fatty acid in *T. jamesii* and both *Asterochloris* species, highlighting a unique aspect of their lipid metabolism. Finally, 1-Monopalmitin (glyceryl palmitate) was the most abundant in *T. cretacea*, although the highest overall levels of this compound were detected in *T. lynniae*.

A substantial portion of the detected compounds consisted of organic acids (Figure 2; Supporting Information Table S8). Several of these are key intermediates in central carbon metabolism, while others play secondary or less well-characterized roles. As observed for amino acids, most organic acids remained below the detection limit in *S. symbiontica* and *M. israelensis*, suggesting a generally lower metabolite content in this species for the same biomass. Among the central carbon metabolism intermediates, malic acid (malate) and lactic acid (lactate) were the most abundant across the majority of species. Malate, a tricarboxylic acid (TCA) cycle intermediate, was the most prominent organic acid in the *Trebouxia* species *T. vagua, Trebouxia* sp. *arnoldoi, T. cretacea, T. corticola* and *T. impressa.* Notably, malate accumulation was significatively higher in species with high ribitol production. In contrast, lactate was the dominant organic acid in *Trebouxia* sp. OTU A25, *T. decolorans*, *M. israelensis*, *S. symbiontica*, *Coccomyxa arvernensis*, *Coenochloris* sp., and *Diplosphaera chodatii*. Gluconic acid emerged as the most abundant organic acid in both *Watanabea* species and *C. vulgaris.* Interestingly, benzoic acid, which does not participate in central carbon metabolism, was the most abundant organic acid in *T. lynniae* and *A. glomerata*, yet undetectable in most other species. This compound is a known precursor to a variety of important plant secondary metabolites. Other prominent organic acids detected included 2,3,4-trihydroxybutanoic acid, the most abundant in *A. erici*, threonic acid, a sugar acid derived from threose, which was most abundant in *T. jamesii*, and 2,3-dihydroxy-2-methylpropanoic acid, a dihydroxy monocarboxylic acid structurally related to propionic acid, found at the highest levels in *V. guanchorum*.

Variable levels of sterols were also detected in the microalgae (Figure 2; Supporting Information Table S9). *D. chodatii* emerged as the most prolific sterol producer, closely followed by *Trebouxia* sp. OTU A25. Campesterol was the predominant sterol in most species, including all *Trebouxia* species, except *Trebouxia* sp. OTU A25, *Trebouxia* sp. *arnoldoi* and *T. impressa,* as well as both *Asterochloris* species, *W. lichenicola, M. israelensis*, *D. chodatii*, and *Coenochloris* sp. In contrast, stigmasterol dominated the sterol profiles of the remaining analyzed species, with the notable exception of *C. arvernensis*, which primarily accumulated β-sitosterol. Additionally, three less abundant sterols were identified: ergosta-4,6,22-triene in *T. cretacea* and *D. chodatii*, and 5ξ-ergost-7-en-3β-ol and ergosterol, detected exclusively in *C. vulgaris*.

Among the significantly different metabolites, phosphoric acid and α-tocopherol were notably widespread across the analyzed species (Figure 2; Supporting Information Table S10). Phosphoric acid, a key molecule involved in energy transfer and metabolic regulation, was detected in all species except *Coenochloris* sp. and *S. symbiontica*, with the highest abundance observed in *Trebouxia* sp. OTU A25. α-Tocopherol, the most biologically active form of vitamin E, was most abundant in *T. asymmetrica*.

Regardless of their specific functional categories, certain metabolites exhibited correlated abundance patterns, suggesting coordinated regulation of their biosynthesis and/or accumulation in microalgae. Supporting Information Fig. S3 illustrates hierarchical clustering of the metabolites most consistently detected across all analyzed species. Notably, the central cluster stands out, comprising 20 metabolites with closely correlated abundance profiles, ranging from amino acids, such as threonine and isoleucine, to monosaccharides, like sorbopyranose and tagatose, and including the most abundant compound, ribitol.

Examining metabolic differences among the studied species reveals two distinct metabolic groups: those that produce and accumulate significant amounts of ribitol, and those that produce negligible amounts of this sugar alcohol, directing carbon toward non-alcohol sugars. This distinction highlights fundamental differences in their metabolic profiles, suggesting divergent biochemical strategies or adaptations across species (Figure 3). These metabolic variations likely reflect underlying ecological pressures, physiological roles, and evolutionary diversification of chemical traits. In analyzing overall metabolic profiles, we observed that metabolites involved in pentose interconversion pathways were notably enriched in species characterized by high ribitol production. However, pathways less directly related to ribitol synthesis also exhibited significant differences. Specifically, among central carbon metabolites, only (S)-malate and D-glucono-1,5-lactone showed significantly higher abundances in high ribitol-producing species compared to others. Additionally, key metabolites for nucleotide sugar synthesis, such as N-acetyl-D-glucosamine and myo-inositol, were overrepresented in these species. In amino acid metabolism, L-proline and (R)-2-hydroxyglutarate similarly showed significantly elevated levels. Within lipid metabolism, ethanolamine phosphate and glycerophosphoglycerol were distinctly enriched in high ribitol producers. Moreover, various metabolites with roles in known metabolic pathways, such as 4-hydroxybutanoic acid, 1,3-propanediol, turanose, allulose, and β-gentiobiose, were also differentially overrepresented in ribitol-rich species. Figure 3 further summarizes metabolites enriched in species with low ribitol accumulation, highlighting distinct metabolic adaptations within this group.

### Exometabolome profiles

The metabolites excreted by selected microalgal species *A. erici*, *T. jamesii*, *W. lichenicola*, and *C. vulgaris* were also analyzed by GC-MS from culture media samples collected during the exponential growth phase. A total of 83 distinct metabolites were identified (Supporting Information Table S11), 72 of which increase or appear during the cultivation of microalgae. Among the species, *T. jamesii* exported the highest number of metabolites (52), followed by *W. lichenicola* (49), *C. vulgaris* (44), and *A. erici* (36). Twenty-five compounds were common across all four species. However, each microalga exhibited distinctive levels of characteristic metabolites (Figure 4a).

**Figure 4.**
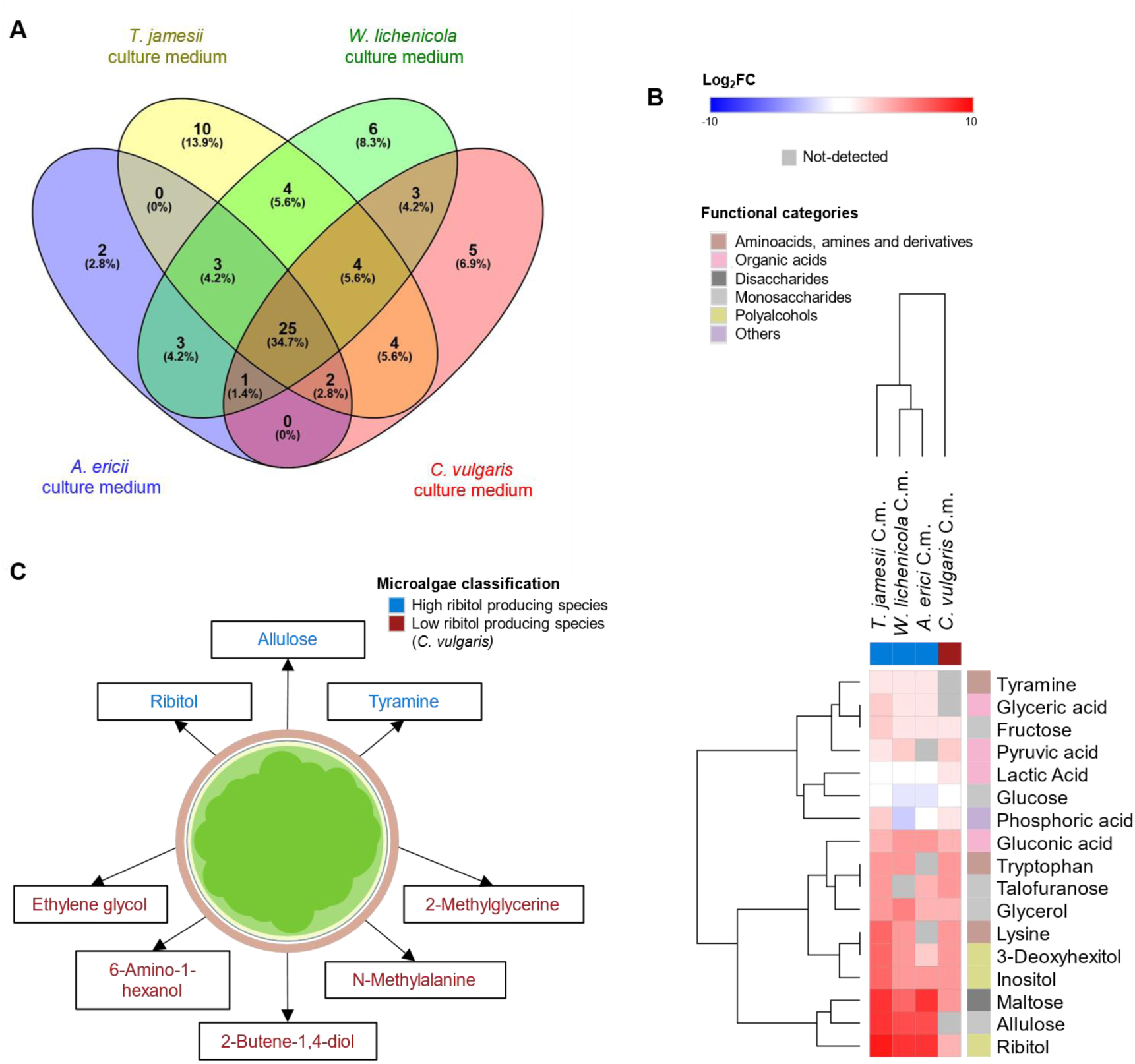
Comparative analysis of metabolites in the culture media (C.m.) of *A. erici*, *T. jamesii*, *W. lichenicola*, and *C. vulgaris* during the late exponential growth phase. a) Venn diagram showing common and unique compounds between microalgae culture media. b) Heatmap illustrating the 17 most represented and significatively different metabolites. Fold-change relative to the blank medium was calculated and log_2_-transformed. Statistical significance was assessed by one-way ANOVA (p-value < 0.05). c) Significatively different metabolites exported by the high and low-ribitol producing species tested. Statistical significance was assessed by Linear mixed-effects (LMM) model (p-value < 0.05).

Notably, *T. jamesii* showed the most diverse excretion profile, with 10 metabolites uniquely identified. As expected, the diversity of excreted compounds was much lower than the diversity of intracellular metabolites detected in the same species (Figure 2). This experimental approach did not reveal a higher diversity of excreted compounds in microalgae capable of forming lichen symbiosis compared to the free-living species. However, the exported metabolome profiles of the symbiotic microalgae were more similar to each other than to the profile of *Chlorella vulgaris* (Figure 4b).

A total of 38 compounds exhibited significant differences between the exported metabolome profiles, with varying proportions among the microalgae, as illustrated in the hierarchical clustered heatmap of the most abundant compounds (Figure 4b). Sugars and sugar alcohols were the most prominently excreted compounds detected. Specifically, ribitol was the most abundant excreted metabolite in *T. jamesii* and *W. lichenicola*, while *A. erici* primarily exported maltose, closely followed by ribitol. In contrast, *C. vulgaris* mainly excreted fructofuranose. Notably, microalgae that produce high endogenous levels of ribitol also export large amounts of this polyol in axenic cultures. The amount of ribitol detected per 100 mg of dry microalgal biomass was approximately equivalent to that present in 6 ml of culture medium in *T. jamesii,* in 3 ml of *W. lichenicola*, or 7 ml of *A. erici* under these experimental conditions. This export pattern extended in part to sugars maltose and allulose, which are also exported in significantly higher amounts by the high-ribitol-producing species than by *C. vulgaris*. Interestingly, *W. lichenicola* exhibited the highest intracellular maltose levels among the analyzed species, yet exported the lowest amount of this sugar among the three symbiotic microalgae. The polyols 3-deoxyhexitol, inositol, and glycerol were excreted differentially by all microalgae, with *T. jamesii* exhibiting the highest levels of the first two and *W. lichenicola* exhibiting the highest levels of glycerol in the culture medium. All four species consumed glucose from the culture medium in differing proportions. *A. erici* and *W. lichenicola* showed the highest glucose uptake under these culture conditions. Additionally, low levels of fructose were detected in the culture media of all four species, and talofuranose was detected in all microalgae except *W. lichenicola*. Notably, sucrose whose endogenous levels are really high in the four species is only exported by *A. erici* and *W. lichenicola*.

Organic nitrogen compounds were present in the culture media at different levels in comparison with the blank medium. Notably, tyramine was exported by all lichen-forming microalgae, while the amino acids tryptophan and lysine were detected in higher levels than in the blank media in all species except *A. erici*. Organic acids were also part of the exported metabolome. Gluconic and lactic acids were exported by all the analyzed microalgae species, as well as glyceric acid. Pyruvic acid in all species except *A. erici*. Finally, phosphoric acid, which is supplied to the medium through its salt forms, showed differential depletion across cultures. *A. erici* and *W. lichenicola* exhibited the greatest uptake, paralleling their high glucose consumption.

Taken together, these results allow us to draw general conclusions about microalgae that produce and export high amounts of ribitol in axenic in vitro cultures, in contrast to *C. vulgaris*, which displays a markedly different metabolomic profile. Several compounds were differentially exported between the high and low-ribitol producing species (Figure 4c). *T. jamesii*, *W. lichenicola*, and *A. erici* export alongside the pentitol ribitol, significantly higher levels of the monosaccharide allulose and tyramine. In contrast, *C. vulgaris* was characterized by the export of the diols ethylene glycol and 2-butane-1,4-diol, the triol 2-methylglycerine, as well as the amino alcohol 6-amino-1-hexanol, and the amino acid derivative, N-methylalanine.

### Implications in the lichen: Ramalina farinacea and Trebouxia jamesii

Metabolomic analyses were performed on the model lichen *Ramalina farinacea*, collected from the bark of *Pinus sylvestris*, and compared with the metabolome of its phycobiont, *T. jamesii*, cultured axenically in vitro. This comparison provides insight into the metabolic behavior of *T. jamesii*, a microalga with high ribitol production (Figure 2) and exportation (Figure 4b) under in vitro conditions, within the context of its symbiotic association in the lichen. A total of 82 distinct metabolites were identified (Supporting Information Table S12). Of these, 28 were unique to the isolated microalga, 32 were exclusive to the lichen, and 22 were shared between both the lichen and its phycobiont (Figure 5b). More than half of the metabolites quantified (50) showed significant differences between the organisms (Figure 5a).

**Figure 5.**
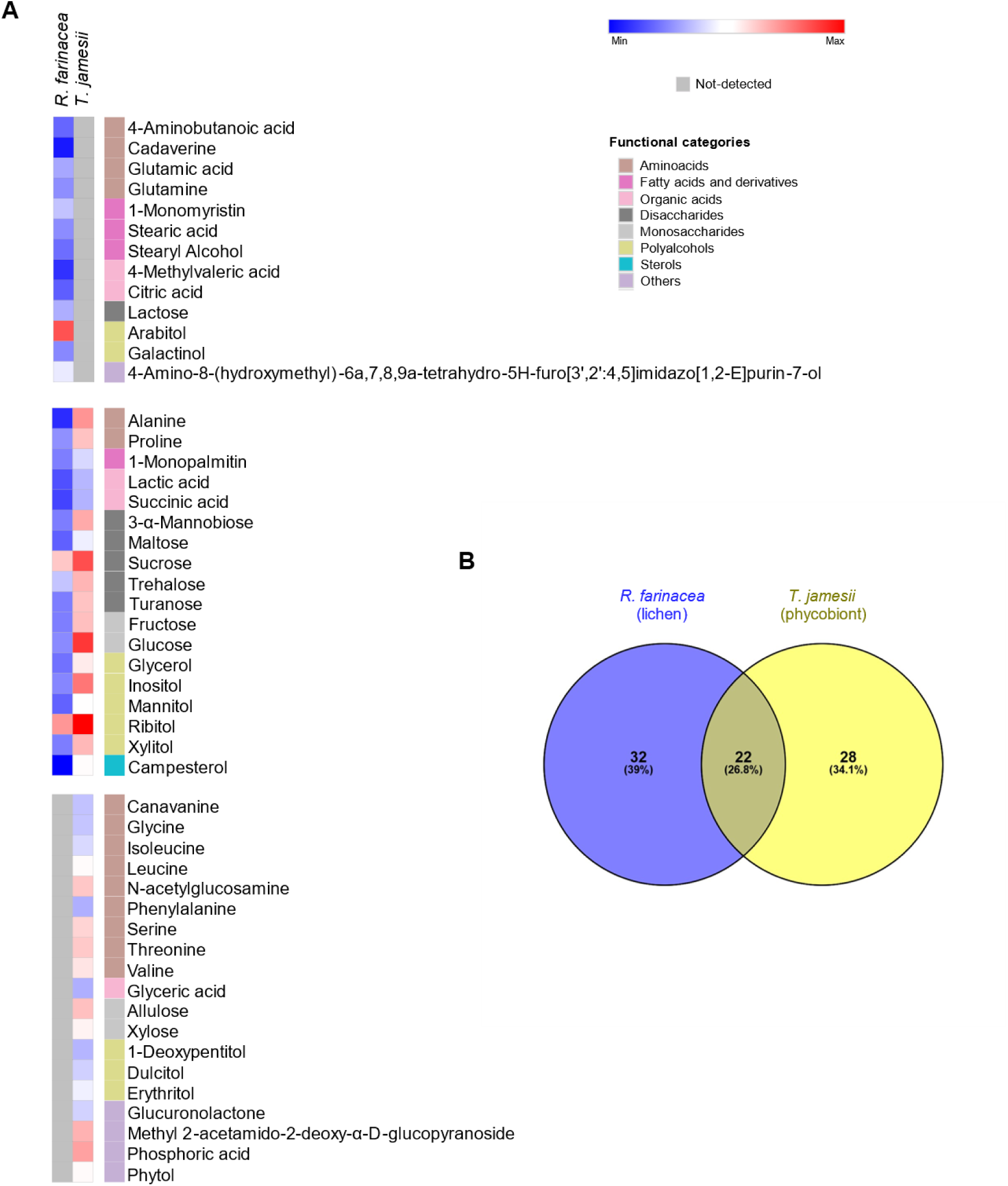
Comparative analysis between *Ramalina farinacea* lichen, and the main phycobiont, *Trebouxia jamesii*. a) Heatmap of the 50 significantly different metabolites between samples. One-way ANOVA analyses were performed for each compound (significance level of p-value < 0.05). Data were log_2_-transformed, and compounds are grouped in functional groups. b) Venn diagram showing common and unique compounds between the lichen and the isolated microalga. Alt text: Graphs labelled A and B. A represents a heatmap of the significant *R. farinacea* and *T. jamesii* metabolites, including functional categories, and hierarchical clustering. B represents a Venn diagram of the *R. farinacea* and *T. jamesii* metabolites.

Variable levels of sugars and sugar alcohols were detected across the analyzed metabolomic profiles. Arabitol was the most abundant compound detected in the metabolome of *R. farinacea*, followed by its stereoisomer ribitol, the most abundant *T. jamesii* compound. While arabitol is not detected in the isolated microalga, ribitol is fifteen times more abundant by total weight in *T. jamesii* in axenic culture than in the collected lichen. Other sugar alcohols present in both organisms include glycerol, inositol, mannitol, and xylitol, all of which were significatively more abundant in the phycobiont. In contrast, the polyol galactinol was detected exclusively in the whole lichen, while 1-deoxypentitol, dulcitol, and erythitol were found only in the isolated phycobiont. Sorbitol and threitol are present in both the lichen and the isolated microalga at comparable levels. Overall, the estimated total content of sugar alcohols in *T. jamesii* was six times higher than in *R. farinacea*. However, sugar alcohols account for 47% of the total quantified metabolites in *T. jamesii*, whereas in *R. farinacea* they represent 77% of the metabolome.

Glucose is the most abundant monosaccharide in the phycobiont and the second most abundant in the lichen, although its level is approximately 300 times higher in the in vitro cultured alga than in the collected lichen, accounting for 13% and 0.3% of the total amount of detected compounds, respectively. The most abundant monosaccharide in *R. farinacea* is tagatose, which is three times more abundant than glucose, and not detected in its phycobiont. Other notable monosaccharides include fructose, which is significantly more abundant in the cultured microalga, as well as allulose and xylose, which are only detected in the microalga and not in the lichen.

Regarding disaccharides, sucrose is the most abundant in both *R. farinacea* and *T. jamesii,* being nine times more abundant in the isolated microalga than in the collected lichen, accounting for 9% and 7.5% of the total amount of detected compounds, respectively. Other disaccharides that are consistently present in higher levels in the cultured phycobiont are 3-α-mannobiose, maltose, trehalose, and turanose. Lactose, however, was only detected in the lichen.

Differential levels of free amino acids and related nitrogenous compounds were also detected in both *R. farinacea* and *T. jamesii.* Alanine was the most abundant amino acid in the cultured phycobiont, whereas glutamic acid predominated in the lichen. As with sugars and their derivatives, overall lower levels of these compounds were substantially lower in the lichen than in the algae. Proline and alanine were present in both organisms, with both being significantly higher in *T. jamesii*. Amino acids glutamic acid, glutamine, and GABA, together with the aliphatic diamine cadaverine, were detected only in the lichen. By contrast, canavanine, glycine, isoleucine, leucine, phenylalanine, serine, threonine, and valine were detected only in *T. jamesii*, together with the amino sugar N-acetylglucosamine.

Fatty acids and their derivative compounds also differed between the lichen and the microalga. Notably, 1-monopalmitin, a monoacylglycerol derived from palmitic acid, was detected in both organisms but was significantly more abundant in *T. jamesii*. In contrast, 1-monomyristin, a monoacylglycerol (and myristic acid derivative), along with stearic acid and its reduced derivative stearyl alcohol, were detected only in the collected *R. farinacea*.

The organic acids profiles of the lichen and the microalga also showed clear differences. Lactic and succinic acids were detected in both organisms, with higher abundance in *T. jamesii*. Glyceric acid was found exclusively in the microalga, whereas citric acid and 4-methylvaleric acid were unique to the lichen *R. farinacea*.

Finally, campesterol, a phytosterol, was present in both organisms but was more prevalent in the phycobiont. Another relevant metabolite is the compound 4-Amino-8-(hydroxymethyl)-6a,7,8,9a-tetrahydro-5H-furo[3’,2’:4,5]imidazo[1,2-E]purin-7-ol, a complex purine derivative unique to the lichen. Notably, methyl 2-acetamido-2-deoxy-α-D-glucopyranoside and phosphoric acid were abundant only in the cultured phycobiont, along with glucuronolactone and phytol.

## Discussion

The analysis of metabolic profiles by using an untargeted metabolomics approach revealed a broad and taxonomically structured biochemical diversity among the studied microalgal species, highlighting both conserved metabolic pathway preferences and unique species-specific adaptations (Figure 1). The variation patterns suggest significant evolutionary divergence among these algae, potentially driven by ecological niche specialization and symbiotic interactions, as suggested by Pichler et al., 2023.

Notably, sugars and sugar alcohols dominated the metabolomic profiles, with ribitol emerging as a pivotal compound in distinguishing metabolite-defined groups (Figure 1b). This sugar alcohol was exported into the culture medium by all the axenically grown lichen microalgal species analyzed (Figure 4b), and was also the second predominant metabolite in the lichen *Ramalina farinacea*, preceded by its stereoisomer arabitol (Figure 5a). The predominance of arabitol in *Ramalina farinacea* aligns with the arabitol accumulation reported in *Ramalina yasudae* (Kosugi *et al*., 2013) and is consistent with the carbon exchange model proposed in other lichen symbioses (Spribille et al. 2022), where the photobiont, here *T. jamesii*, delivers a photosynthesis-derived sugar alcohol, ribitol, to *R. farinacea* mycobiont, which converts and stores it as arabitol.

Ribitol production correlated broadly with phylogenetic affiliations in microalgae, suggesting chemotaxonomic relevance (Figure 1c). However, notable exceptions, such as *W. acidophila* and *W. lichenicola*, emphasize the complex interplay between phylogeny, ecological roles, and metabolic evolution (Gustavs et al., 2011; Hotter et al., 2018). Ribitol’s dominance in the metabolome of most analyzed lichen-forming microalgae highlights its role in lichen symbiosis, facilitating symbiotic stability through carbon exchange and osmoregulation (Wang *et al*., 2009; Meeßen *et al*., 2013; Scharnagl *et al*., 2023). However, species such as *Diplosphaera chodatii* and *Coccomyxa arvernensis*, despite their presence within lichen symbioses, exhibit metabolomic profiles more similar to the analyzed free-living algae (Figure 2, Supporting Information Fig. S3a). They showed limited ribitol accumulation, coupled with comparatively higher levels of sorbitol, a polyol not dominant in their overall metabolomes. Sorbitol production has been previously reported in *D. chodatii* (Wang *et al*., 2014; Medwed *et al*., 2021; Nazem-Bokaee *et al*., 2025) but not yet documented in *C. arvernensis*, which is described to produce moderate ribitol levels by (Gustavs et al., 2011). The predominance of sorbitol and the reduced abundance of other polyalcohols indicate that these two species adopt a distinct biochemical strategy, suggesting they may fulfill a different role within the lichen in terms of metabolic relationship, which could have physiological and ecological implications. On the other hand, the aeroterrestrial species *Coenochloris sp.*, which is not known to engage in lichen symbioses, also produces glycerol but in lower amounts than most of the symbiotic microalgae, having a much lower sugar alcohol content, whereas the aquatic free-living species *Chlorella vulgaris* produces negligible levels of sugar alcohols (Figure 2). Overall, algae species that accumulate sugar alcohols as their predominant low-molecular-weight metabolites under controlled, favorable conditions are more frequently associated in lichen symbioses (Sanders & Masumoto, 2021). In particular, taxa that produce ribitol as their main polyol tend to form the vast majority of lichen associations, in contrast to those that primarily synthesize sorbitol or other sugar alcohols (Figure 1c). Together, the genera *Trebouxia* and *Asterochloris*, and to a lesser extent *Myrmecia*, *Symbiochloris*, *Vulcanochloris*, and *Watanabea*, serve as phycobionts in over ten times more lichen partnerships than *Diplosphaera* or *Coccomyxa* species. In contrast, microalgae such as *Coenochloris* sp. and *Chlorella vulgaris*, which produce minimal sugar alcohols, are not observed as lichen phycobionts. Collectively, these patterns suggest that high ribitol production and export may represent a preferential biochemical strategy in lichen symbiosis.

Similarly, these findings suggest that the gradient of constitutive sugar alcohol accumulation observed under favorable culture conditions may serve as a biochemical marker of each microalga’s degree of desiccation tolerance and ecological adaptation in its natural environment (Gasulla *et al*., 2021). The differential co-accumulation of sugar alcohols, sugars, amino acids, and organic acids detected in this study aligns with their roles as compatible solutes or osmoprotectants, a well-established mechanism in plant cells to maintain cellular homeostasis under abiotic stress (Mehta & Vyas, 2023). Polyols in particular, which were among the most abundant and variable metabolites across the studied species, are recognized for their roles in stabilizing proteins, preserving membrane integrity, and mitigating oxidative damage during desiccation and osmotic shock (Gustavs *et al*., 2010; McParland *et al*., 2021). Their accumulation in higher quantities in lichen-forming microalgae likely contributes to the remarkable desiccation tolerance characteristic of symbiotic phycobionts, which is directly correlated with tolerance to other abiotic stresses. Beyond their classical role as osmoprotectants, many of the highly accumulated metabolites in the high ribitol-producing microalgae, such as glycerol, inositol, proline, and malate, are potential components of Natural Deep Eutectic Solvents (NADES). These mixtures form viscous, stable liquid phases even in low-water or desiccated states, offering protection to macromolecules and maintaining enzymatic activity under extreme conditions (Cajnko *et al*., 2023). This facilitates both the protection of macromolecules during periodic desiccation and the rapid reactivation of metabolic processes upon rehydration. It should be noted that sucrose is also present in large quantities across all analyzed *Trebouxiophyceae* species, whether lichen-associated or free-living (Figure 2), and it also contributes to osmoprotection and stress mitigation by the same mechanisms. However, species that accumulate ribitol may have an added advantage, as polyols possess a greater capacity for hydrogen bonding, thereby enhancing their ability to stabilize macromolecules under conditions of desiccation or osmotic stress. The importance of ribitol and sucrose is evident in *R. farinacea* and its phycobiont *T. jamesii*, as these are the two metabolites consistently present at high levels in their metabolomes (Fig. 5a). Ribitol was found to be four times more abundant in the alga grown in a nutrient-rich medium than in the whole lichen collected in the field, sucrose was nine times more abundant, and glucose was nearly 300 times higher (Supporting Information Table S12). This pattern suggests that, within the lichen, the alga actively prioritizes the maintenance of elevated ribitol levels, together with sucrose, to a lesser extent, even under suboptimal conditions. In contrast, common monosaccharides such as glucose appear to be accumulated mainly when resources are abundant or under free-living conditions, indicating a metabolic shift in the symbiotic state toward specialized carbohydrate allocation.

The combined intra- and extracellular metabolomic profiles of each species are critical both for determining their capacity to establish specific lichen symbioses and for enabling adaptation to the environmental conditions of their natural habitats.

This study reveals that maltose and allulose exhibit an exportation pattern in the analyzed lichen-forming microalgae that closely parallels that of ribitol (Figure 4c), a novel observation not previously reported in the literature. However, both sugars are present at much lower levels than ribitol and are less prominent within the overall metabolome. Notably, while maltose is consistently detected both intracellularly (Figure 2) and extracellularly (Figure 5a), its relative abundance is modest. In contrast, allulose is not detected endogenously in some of the high ribitol-producing species (Figure 2), despite being present in their exometabolome (Figure 5a). A similar trend is observed in *Ramalina farinacea*, where maltose is present, but allulose is not detected (Figure 5a). These findings suggest that although maltose and allulose may be co-exported with ribitol, their roles in lichen symbiosis may differ and they could be secondary to that of ribitol. In contrast, glucose, and sucrose, the other two generally dominant sugars within intracellular metabolomes, show different patterns of export in axenic phycobiont cultures. Glucose is consumed rather than exported by all studied microalgae species, resulting in an overall net uptake from the culture medium. Sucrose, however, is actively exported by *A. erici* and *W. lichenicola* but not by *T. jamesii* (Figure 5a). These findings contrast with the observations reported in *D. chodatii,* which releases sorbitol, glucose, and sucrose that are incorporated by the mycobiont when co-cultivated, but not in axenic cultures (Wang *et al*., 2014). The differential exportation patterns between lichen microalgae suggest distinct metabolic strategies and roles for these sugars within each phycobiont and their lichen symbioses. In addition, rare sugars β-gentiobiose and turanose, and polyols glycerol and inositol are produced in higher levels by high ribitol-producing microalgae (Figure 2), some of them being differentially exported by the lichen microalgae (Figure 5a). Their chemical properties suggest a potential function as compatible osmolytes, although they are also likely to possess additional unexplored roles. Furthermore, in high ribitol producing species, the consistently elevated malate levels point to an enhanced TCA cycle flux feeding polyol biosynthesis (Fig. 3). Acting as a central metabolic hub, malate also supports photosynthetic redox balance via the malate valve and links carbon metabolism to nitrogen assimilation and photorespiration, thereby maintaining metabolic flexibility under variable environmental conditions. This integration of energy, carbon, and redox management (Dao *et al*., 2022) likely provides a physiological advantage in lichen symbiosis. Moreover, malate’s osmoprotective capacity and potential role as a Natural Deep Eutectic Solvent (NADES) component (Cajnko *et al*., 2023) further strengthen stress resilience, reinforcing the adaptive advantage of these microalgae in desiccation-prone habitats.

In conjunction with these findings, the widespread accumulation of the non-proteinogenic amino acid pyroglutamic acid (5-oxoproline) in all the analyzed species, and especially its high abundance in the high ribitol-producing species (Figure 2), could be related to its proposed role as an analogue or reservoir of glutamate, as well as for osmoprotection and antioxidant activity, even though its role had not been previously reported in green microalgae (Kumar *et al*., 2012). High ribitol producers also constitutively accumulate proline (Figure 2 and Figure 3), a renowned compatible osmolyte and a classical stress-responsive amino acid in plants and microalgae, indicating a permanently primed stress-tolerant state (Barera & Forlani, 2023). They do it together with the constitutive accumulation of 2-hydroxyglutarate (Figure 3), which is reported to accumulate in the model green microalgae *C. reinhardtii* as a signaling molecule for osmotic stress (Colina *et al*., 2020). In contrast, the preferential accumulation of GABA in all low ribitol-producing species except *C. vulgaris* (Figure 3) points to an alternative amino acid based strategy in these three species, potentially linked to their abiotic stress response and lipid metabolism(Zhao *et al*., 2020; Li *et al*., 2021). The clear partitioning of free amino acid accumulation patterns between high and low ribitol producers highlights the existence of divergent biochemical strategies for osmotic adaptation in the analyzed microalgae, also different from other microalgae in the literature (León-Vaz *et al*., 2023). Moreover, the differential excretion of amino acid derivative, tyramine, predominantly by lichen-forming microalgae and closely paralleling the export pattern of ribitol (Figure 4b), while at much lower concentrations, may indicate roles in symbiotic or environmental communication. The excretion of these compounds has not previously been reported in lichen microalgae and indicates potential paths for further ecological and functional studies.

Phosphate-containing compounds involved in lipid metabolism also emerge as key differentiators between the analyzed groups of microalgae. Glycerophosphoglycerol and ethanolamine phosphate were significantly more abundant in high ribitol-producing species (Figure 3), yet their occurrence and function in microalgae remain largely unexplored. In other systems, glycerophosphoglycerol is recognized as a membrane phospholipid that contributes to membrane structure and dynamics (Trötzmüller, 2016), while ethanolamine phosphate (phosphoethanolamine) is a well-established intermediate in the phosphatidylethanolamine biosynthesis pathway, a major route for membrane lipid production (Hanchi *et al*., 2018). These known roles suggest that, in microalgae, both compounds may contribute differentially to the maintenance and adaptive modulation of membrane integrity in high ribitol-producing species.

It is worth noting the accumulation and export pattern of inorganic phosphorus between lichen symbionts. *T. jamesii* accumulates high intracellular levels of phosphoric acid (Figure 2) and releases it into the medium under axenic culture conditions. In contrast, in the *R. farinacea* symbiosis, phosphoric acid is not detected at appreciable levels in the lichen thallus, despite its presence in the isolated phycobiont. This pattern suggests potential transfer and metabolic utilization of phosphoric acid within the symbiotic association. Interestingly, this contrasts with observations regarding phosphoric acid in resynthesized *Usnea hakonensis* thalli (Kono *et al*., 2020), where inorganic phosphate transport is proposed to occur predominantly from the mycobiont to the photobiont. Our results suggest that in *R. farinacea*, inorganic phosphate transfer may not be strictly unidirectional and could occur in the opposite direction in phosphoric acid form, from the phycobiont to the mycobiont.

Altogether, our findings propose that metabolic features such as sugar alcohol composition, osmoprotectant accumulation, and selective metabolite accumulation and excretion are part of the mechanisms that support lichen symbiosis. Particularly, ribitol may be a pivotal biochemical trait driving symbiotic capability in Trebouxiophyceae. These insights contribute to our understanding of the biochemical determinants of symbiotic compatibility and offer new perspectives for using metabolomics in chemotaxonomic classification and evolutionary ecology that could be applied to predict symbiotic compatibility in untested algal-fungal pairs. This study provides a foundational metabolomic framework to identify key traits associated with lichenization and guides future research into metabolic signaling and symbiotic specificity.

## Acknowledgments

This study was supported by projects PROMETEO/2021/005, PROMETEO2024-044, AGROALNEXT2022-061 (Generalitat Valenciana), and PID2021-127087NB-I00 (Ministerio de Ciencia e Innovación) awarded to P.C. and the predoctoral project ESA Idea I-2021-05232 (European Space Agency) awarded to M.P.-R. We thank Dr Sales Galletero and Isabel Lliso (SCSIE UV) for their assistance with mass spectrometry analyses.

## Author contributions

Design of the research, M.P.-R.; performance of the research, M.P.-R. and D.C.; data analysis, M.P.-R.; writing-original draft preparation, M.P.-R.; writing-review, all authors; project administration, P.C.; funding acquisition, P.C., F.M., and M.P.-R. All authors have read and agreed to the published version of the manuscript.

## Data availability

The data is available within the article and its supplementary materials.

## Supporting Information

Additional Supporting Information may be found online in the Supporting Information section at the end of the article.

**Supplementary Table 1.** Average normalized peak areas for each metabolite of the 21 microalgae species analyzed.

**Supplementary Table 2.** UpSet diagram datasets.

**Supplementary Table 3.** Average normalized peak areas of the five most abundant metabolites detected in each of the 21 microalgal species analyzed.

**Supplementary Table 4.** Sugar alcohols average normalized areas for each species.

**Supplementary Table 5.** Sugars average normalized areas for each species.

**Supplementary Table 6.** Amino acids average normalized areas for each species.

**Supplementary Table 7.** Fatty acids average normalized areas for each species.

**Supplementary Table 8.** Organic acids average normalized areas for each species.

**Supplementary Table 9.** Sterols average normalized areas for each species.

**Supplementary Table 10.** Other significant compounds average normalized areas for each species.

**Supplementary Table 11.** Log_2_ fold-change of the average normalized peak areas for the metabolites detected in microalgal culture media compared to the blank medium control.

**Supplementary Table 12.** Average normalized peak areas for each metabolite of *R. farinacea*and *T. jamesii*.

**Supplementary Figure 1.** Principal Component Analysis (PCA) of the metabolomic profiles of the 21 microalgal species analyzed. Metabolite data were obtained via GC-MS, with three biological replicates and two technical replicates per species. The solid black line separates species predominantly producing ribitol (right) from those in which sucrose or glucose is the major metabolite (left).

**Supplementary Figure 2.** Relative abundance of the top five most abundant metabolites per microalgal species. Stacked bar plots represent the total metabolite profile per species, normalized to 100%. The five most abundant metabolites per species are shown in colors, while all other detected compounds are grouped into the "Others" category (grey).

**Supplementary Figure 3.** Similarity matrices illustrating relationships among a) the analyzed species based on their complete metabolomic profiles and b) the 48 most significantly represented metabolites across species, grouped by functional categories. Clustering was performed using Pearson correlation coefficients. Statistical significance for each metabolite was determined by one-way ANOVA (p-value < 0.05).

